# *FLCN* Gene Ablation Reduces Fibrosis and Inflammation in a Diet-Induced NASH Model

**DOI:** 10.1101/2020.09.10.291617

**Authors:** Mathieu Paquette, Ming Yan, Leeanna El-Houjeiri, Marco Biondini, Josué M. J. Ramírez-Reyes, Alain Pacis, Hyeonju Jeong, Jennifer L. Estall, Peter M. Siegel, Arnim Pause

**Affiliations:** Goodman Cancer Research Center, McGill University, Montréal, Québec, Canada; Department of Biochemistry, McGill University, Montréal, Québec, Canada; Canadian Centre for Computational Genomics, McGill Genome Centre, Montréal, Québec, Canada; Institut de recherches cliniques de Montréal (IRCM), Montréal, Québec, Canada; Department of Medicine, McGill University, Montréal, Québec, Canada

**Keywords:** non-alcoholic steatohepatitis (NASH), methionine- and choline-deficient (MCD) diet, fibrosis, inflammation, autophagy, FLCN

## Abstract

Non-alcoholic steatohepatitis (NASH) represents a major economic burden and is characterized by triglyceride accumulation, inflammation, and fibrosis. No pharmacological agents are currently approved to treat this condition. Emerging data suggests an important role of autophagy in this condition, which serves to degrade intracellular lipid stores, reduce hepatocellular damage, and dampen inflammation. Autophagy is primarily regulated by the transcription factors TFEB and TFE3, which are negatively regulated by mTORC1. Given that FLCN is an mTORC1 activator via its GAP activity towards RagC/D, we generated a liver specific *Flcn* knockout mouse model to study its role in NASH progression. We demonstrate that loss of FLCN results in reduced triglyceride accumulation, fibrosis, and inflammation in mice exposed to a NASH-inducing diet. Hence, the GAP activity of FLCN could a promising target for small molecule drugs to treat NASH progression by specifically activating autophagy and lysosomal biogenesis while leaving mRNA translation machinery unperturbed. Collectively, these results show an unexpected role for FLCN in NASH progression and highlight new possibilities for treatment strategies through its role in hepatocyte homeostasis.

## Introduction

Obesity prevalence continues to rise drastically worldwide and represents a major health challenge as it increases the risk of developing diabetes [1], cardiovascular diseases [1,2], poor mental health [3], and several cancers [4]. Worldwide, nearly one third of the population is classified as overweight or obese, a number that doubled since 1980 [5]. Non-alcoholic fatty liver disease (NAFLD) is a disease spectrum characterized by triglyceride accumulation in hepatocytes and is associated with obesity, insulin resistance, type 2 diabetes, and dyslipidemia [6]. Approximately one-third of patients with NAFLD will progress toward non-alcoholic steatohepatitis (NASH) [7]. Furthermore, approximately one-third of patients with NASH will progress to cirrhosis and final liver dysfunction [8,9]. There is currently no pharmacological agent approved to treat NASH or to slow its progression [10]; the only proven treatments are weight loss and increased physical activity [11].

NASH is characterized by hepatocellular injury, immune cell-mediated inflammation, and progressive liver fibrosis [12]. The accumulation of lipid intermediates within hepatocytes causes lipotoxicity, cellular stress, and eventually cell death via distinct mechanisms [12]. Toxicity arises from an increase in reactive oxygen species production due to enhanced fatty acid oxidation [13,14]. Moreover, excess fatty acid modulates cell signaling, disrupts metabolic homeostasis, and induces ER stress [15,16]. Damaged livers constitute a unique pro-inflammatory microenvironment composed of a variety of immunologically active cells, such as Kupffer cells, T cells, and hepatic stellate cells (HSC) [17]. The inflammation observed in NASH livers may be a consequence of lipotoxicity where chronic hepatocyte death generates damage-associated molecular patterns (DAMPs), attracting immune cells in the liver [18]. Yet, a sustained inflammatory response can further contribute to hepatocellular damage, creating a feed-forward loop toward liver injury [12].

Chronic liver injury eventually leads to fibrosis as a consequence of chronic wound-healing responses initiated by dying hepatocytes [19]. In a healthy state, wound-healing restores liver function and structure by recruiting liver progenitor populations and inflammatory cells, which can release angiogenic and matrix remodeling factors [20]. In NASH, wound-healing is perpetual, and the persistent inflammation prevents the termination of the process [19]. This results in excessive liver progenitors and myofibroblast accumulation, scarring (cirrhosis), and matrix remodeling, perpetuating fibrogenesis and inflammatory responses [15].

Emerging data supports a role of autophagy in the liver to regulate intracellular lipid stores and energy homeostasis. Autophagy is an important process in which the cells degrade its own content by sequestering cargoes inside the lysosomes. Numerous studies have reported that obesity impairs autophagy and lysosome function and can lead to hepatocellular injury [21–27]. Moreover, autophagy can reduce inflammation by suppressing transcription or maturation of pro-inflammatory cytokines [28,29]. Degradation of cargoes inside the lysosome is a multistep process that is tightly regulated by multiple regulators known as “autophagy-related proteins”, but also by master transcription factors such as Transcription Factor EB (TFEB) and Transcription Factor E3 (TFE3) [30,31]. The best described regulator of TFEB and TFE3 is the serine/threonine kinase mammalian Target of Rapamycin Complex 1 (mTORC1). mTORC1 regulates cell proliferation and growth by integrating stress and growth signals to promote anabolic processes and inhibit catabolic processes such as autophagy [32]. Under non-stress conditions, mTORC1 directly phosphorylates TFEB and TFE3, resulting in cytoplasmic sequestration and inhibition of their activity [33–36]. Conversely, inhibition of mTORC1 removes the repressive phosphorylation on TFEB and TFE3, resulting in their nuclear translocation and activation of a panel of genes involved in autophagy, lysosomal biogenesis, lipid metabolism, innate immune response, and pathogen resistance [37–44].

We and others have shown that Folliculin (FLCN) is a negative regulator of TFEB and TFE3 and positive regulator of mTORC1 [43,45–47]. Loss-of-function mutations in human FLCN are associated with the Birt-Hogg-Dubé (BHD) syndrome, a disease characterized by skin fibrofolliculoma, lung cysts, increased risk for spontaneous pneumothorax and predisposition to renal cell carcinoma (RCC) [48]. Moreover, FLCN exhibits GTPase-activating protein (GAP) activity upon nutrient replenishment targeting the Ras-related GTPase C and Ras-related GTPase D (RagC/D) resulting in mTORC1 activation, inhibition of TFEB and TFE3 nuclear translocation, and impaired autophagy and lysosomal function [49–51]. Upon nutrient depletion FLCN activity is being inhibited via an unknown sensing mechanism, resulting in inhibition of mTORC1 activity, TFEB and TFE3 activation, and stimulation of autophagy and lysosomal activity [46,52]. Similarly, complete loss of FLCN or FLCN loss-of-function mutations identified in BHD patients phenocopied the effect observed during nutrient depletion [45,46,50].

FLCN loss of function has been studied in in nematodes and various tissues of knockout mice with surprisingly beneficial effects [46,53–56]. FLCN deficient nematodes develop normally and show resistance to oxidative stress, heat, anoxia, hyperosmotic stresses, and to bacterial pathogens [43,53,54]. Mice with a conditional FLCN knockout in skeletal muscle developed a pronounced metabolic shift towards oxidative phosphorylation, increased mitochondrial biogenesis, and red coloured muscles [55]. Conditional FLCN adipocyte knockout mice developed resistance to high-fat diet (HFD)-induced obesity as well as browning of white adipocytes and increased activity of brown fat [46,56]. Little is known about how FLCN regulates a highly metabolic tissue such as the liver. More specifically, the potential role of FLCN in NASH progression has never been investigated. In this study, we generated a liver specific *Flcn* knockout mouse model challenged with a methionine/choline deficient (MCD) diet. We demonstrate that liver-FLCN KO mice were protected from MCD-induced triglyceride accumulation and liver damage. Importantly, loss of FLCN in liver tissue resulted in increased autophagic activity and nuclear localization of TFE3 and TFEB, reduced fibrosis, and reduced inflammation, underlying potential mechanisms for NASH protection.

## Results

### FLCN loss reduces high fat diet induced NAFLD

To understand the role of FLCN in the liver, we generated a liver-specific FLCN KO mouse model by crossing *Flcn*^lox/lox^ mice with albumin-cre^+/-^ mice (Supplemental Figure 1A-C) [56]. Albumin-cre expression is restricted to hepatocytes [57]. Albumin-cre FLCN knockout mice were born at the expected Mendelian frequency, displayed no developmental defects, survived without difficulty, were fertile, and had similar weights compared with wild-type mice at weaning (4 weeks of age). We first established whether FLCN loss in the liver affects whole-body metabolism by feeding them either normal chow or a high fat diet (HFD) containing 60% kcal from fat for 6 weeks. FLCN KO mice displayed resistance to weight gain when maintained on HFD compared to control mice expressing the albumin-cre transgene without floxed alleles (cre) (Supplemental Figure 1D). Hematoxylin and eosin (H&E) staining revealed a strong reduction in fat droplets in livers of FLCN KO mice when challenged with HFD compared to cre mice (Supplemental Figure 1E). FLCN KO mice also exhibited superior glucose handling during both glucose tolerance tests (GTTs) and insulin tolerance tests (ITTs) (Supplemental Figure 1F-G). Hence, loss of FLCN specifically in the liver improved metabolic homeostasis and prevented weight gain in a diet-induced obesity and NAFLD model.

### FLCN loss reduces MCD diet-induced liver triglyceride accumulation and liver damage

Since liver phenotypes associated with high fat feeding alone are highly associated with obesity and whole-body insulin resistance, the effects observed in liver may have been an indirect consequence of reduced adiposity. To test whether protection against steatosis and NAFLD was due to cell-autonomous effects of FLCN KO in hepatocytes, we investigated whether loss of hepatic FLCN was also protective in a model of NASH that does not cause weight gain or insulin resistance. To this end, we subjected cre and liver-FLCN KO mice to a methionine and choline deficient (MCD) diet for six weeks. This diet is a model of a metabolic challenge and results in a significant and rapid onset of the NASH phenotype with fibrosis, inflammation, oxidative stress, and liver cell death [58]. As previously described [59], we show that this diet induced significant weight loss (Supplemental Figure 2A). No weight differences were observed between the two genotypes, both in male and female mice (Supplemental Figure 2A). After six weeks on the MCD diet, mouse serum was extracted from cardiac punctures and livers collected. H&E staining revealed a partial reduction in fat droplets in FLCN KO mice compared to cre mice (Figure 1A). Specific quantification of the total triglycerides in the liver tissue also showed a significant reduction in fat accumulation in FLCN KO livers (Figure 1B). We subsequently assessed multiple blood parameters. Serum total cholesterol, serum free cholesterol, and serum cholesterol esters were all significantly reduced in both male and female mice fed the MCD diet; however, there were no significant differences between cre and FLCN KO mice (Supplemental Figure 2B-D). Similarly, serum triglycerides were not significantly different between both genotypes (Figure 1C). When hepatocytes undergo apoptosis, they release the enzyme alanine transaminase (ALT), which is used as a readout of liver damage. Interestingly, a reduction in ALT levels were observed in FLCN KO mice fed the MCD diet compared to cre (Figure 1D). These results suggest that liver-specific loss of FLCN reduces fat accumulation and liver damage through another mechanism than increased fat export in the blood.

**Figure 1:**
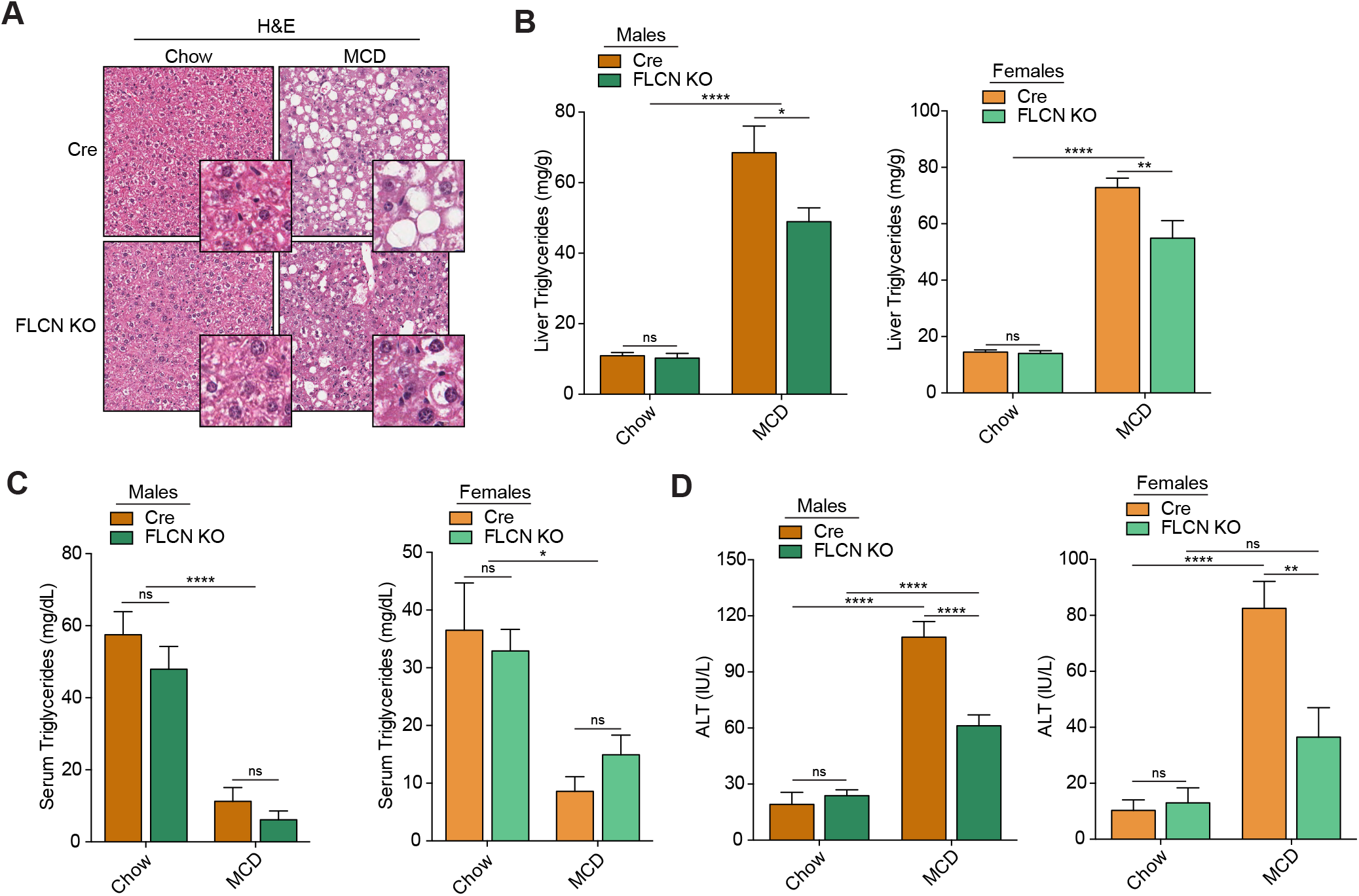
FLCN loss reduces MCD diet induced liver triglyceride accumulation without affecting blood parameters. (**A**) H&E staining in liver sections of mice fed either chow or methionine/choline deficient diet (MCD) over a period of 6 weeks. Images are representative of 8 mice per condition. (**B**) Liver triglycerides quantification in male (left) and female (right) mice fed as described in (A) (mean ± SEM, two-way ANOVA, ns = not significant; *P < 0.05; **P < 0.01; ****P < 0.0001; n = 8 mice per condition). (**C**) Serum triglycerides quantification in male (left) and female (right) mice fed as described in (A) (mean ± SEM, two-way ANOVA, ns = not significant; *P < 0.05; ****P < 0.0001; n = 8 mice per condition). (**D**) Alanine transaminase (ALT) quantification in male (left) and female (right) mice fed as described in (A) (mean ± SEM, two-way ANOVA, ns = not significant; **P < 0.01; ****P < 0.0001; n = 8 mice per condition).

### RNA-seq reveals major pathways affected by FLCN loss

These results prompted us to investigate the molecular signature of FLCN-null livers responsible for the protective phenotype. Total RNA was purified from liver tissue homogenates (6 weeks on diet) and high-throughput whole-transcriptome sequencing (mRNA-seq) performed. Unsupervised hierarchical clustering highlighted three distinct patterns of gene expression changes (Figure 2A, Supplementary table 1). One cluster displayed a clear increase in gene expression induced by the MCD diet, which was not affected by FLCN loss (cluster 1, Figure 2A). Another cluster included genes whose expression was diminished in mice maintained on the MCD diet, which were again unaffected by FLCN status (cluster 3, Figure 2A). A third cluster highlighted genes whose expression was induced in cre mice receiving the MCD diet that were not increased in FLCN KO mice maintained on the MCD diet (cluster 2, Figure 2A-B). Gene enrichment analysis using GO Enrichr revealed the major pathways differentially affected in this cluster by the diet and the genotypes (Figure 2C). The most significantly affected biological pathway was extracellular matrix organization, which represents genes involved in fibrosis development typically observed in NASH. Other important pathways affected include mitotic sister chromatid segregation, sister chromatic segregation, and mitotic nuclear division. These genes are generally induced in regenerating livers following injury [60]. Moreover, pathways involved in inflammation such as neutrophil activation and cellular response to cytokine stimulus were differentially expressed. Concurrently, no significant changes in expression between cre and FLCN KO mice were detected for pathways involved in apoptosis (GO:0006915), lipolysis (GO:0016042), lipogenesis (GO:0008610), glycolysis (GO:0006096), and gluconeogenesis (GO:0006094) (Supplemental figure 3A-E). Relative mRNA transcript levels were also not different between the genotypes (Supplemental figure 3F). Overall, genes involved in fibrosis (GO:0030198), inflammation (GO:0006954) and mitotic cell cycle (GO:0000278) were induced in mice fed the MCD diet but to a lesser extent in FLCN KO mice (Figure 2D-F).

**Figure 2:**
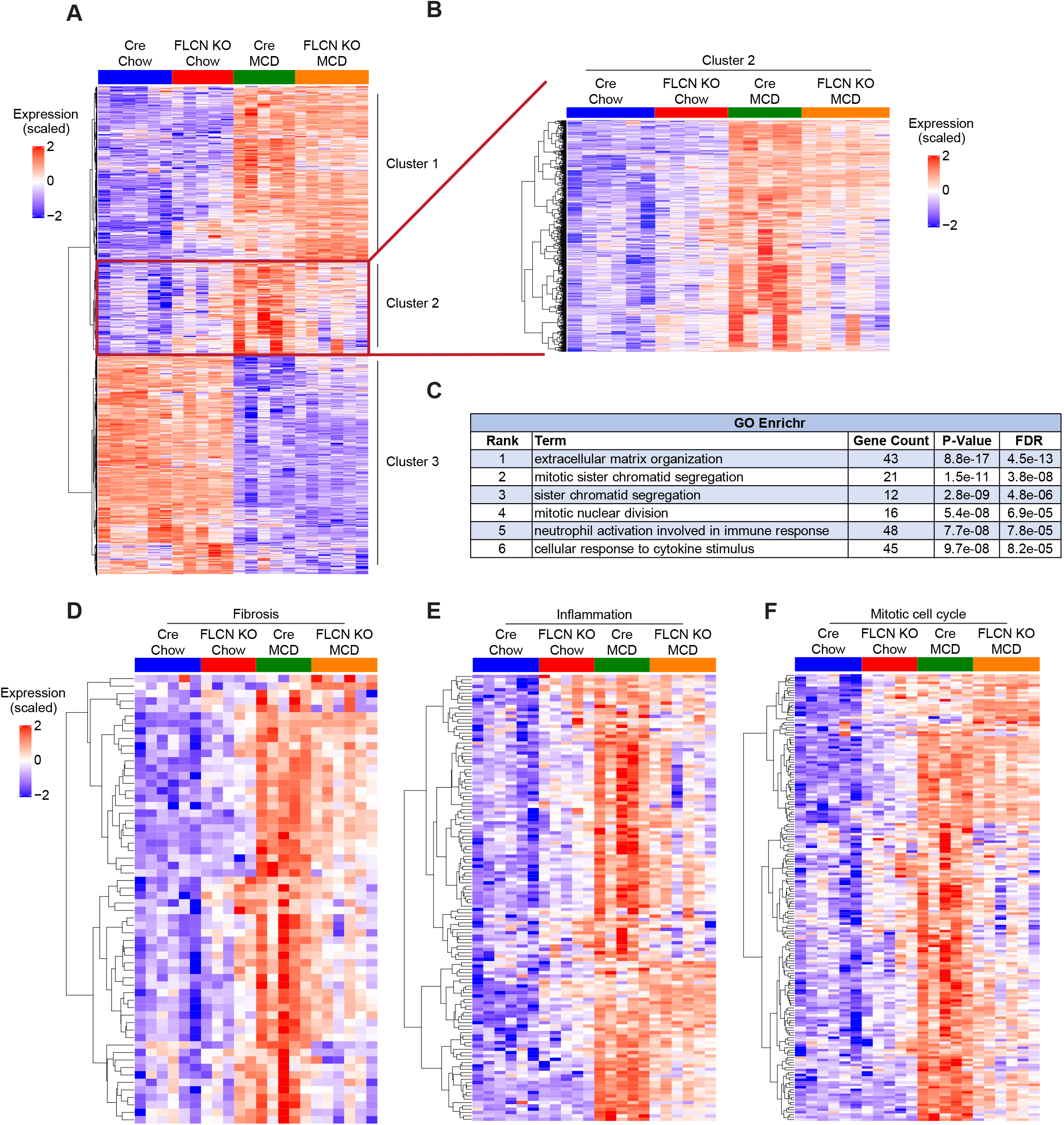
RNA-seq reveals major pathways affected by FLCN loss. (**A-B**) Unsupervised hierarchical clustering following RNA-seq analysis in livers of mice fed either chow or methionine/choline deficient diet (MCD) over a period of 6 weeks. (**C**) Gene enrichment analysis using GO Enrichr. (**D**) Unsupervised hierarchical clustering following RNA-seq analysis of fibrosis-related genes (GO:0030198). (**E**) Unsupervised hierarchical clustering following RNA-seq analysis of inflammation-related genes (GO:0006954). (**F**) Unsupervised hierarchical clustering following RNA-seq analysis of mitotic cell cycle-related genes (GO:0000278).

### FLCN loss reduces MCD diet-induced fibrosis and inflammation

One hallmark of NASH is fibrosis, which generates liver damage and eventually leads to hepatocyte death and liver dysfunction [61]. We validated the RNA-seq data by measuring the relative mRNA transcript levels of genes related to fibrosis; all were induced in cre mice fed the MCD diet, but generally to a lower extent in FLCN KO mice (Figure 3A). Sirius Red staining revealed that fibrosis was aggravated in cre mice challenged on the MCD diet, but this effect was diminished in FLCN KO mice (Figure 3B, quantified in C). Immunohistochemistry staining of 0-smooth muscle actin (⍰-SMA), a fibrosis-related marker, was significantly increased in cre mice challenged with the MCD diet but to a lesser extent in FLCN KO mice (Figure 3D, quantified in E). Finally, immunohistochemistry staining of 4-Hydroxynonenal (4-HNE), a lipid peroxidation marker, was significantly increased in cre mice challenged with the MCD diet but diminished in FLCN KO mice (Figure 3F, quantified in G). Taken together, these results demonstrate that liver-specific loss of FLCN reduces fibrosis, protecting from diet-induced NASH.

**Figure 3:**
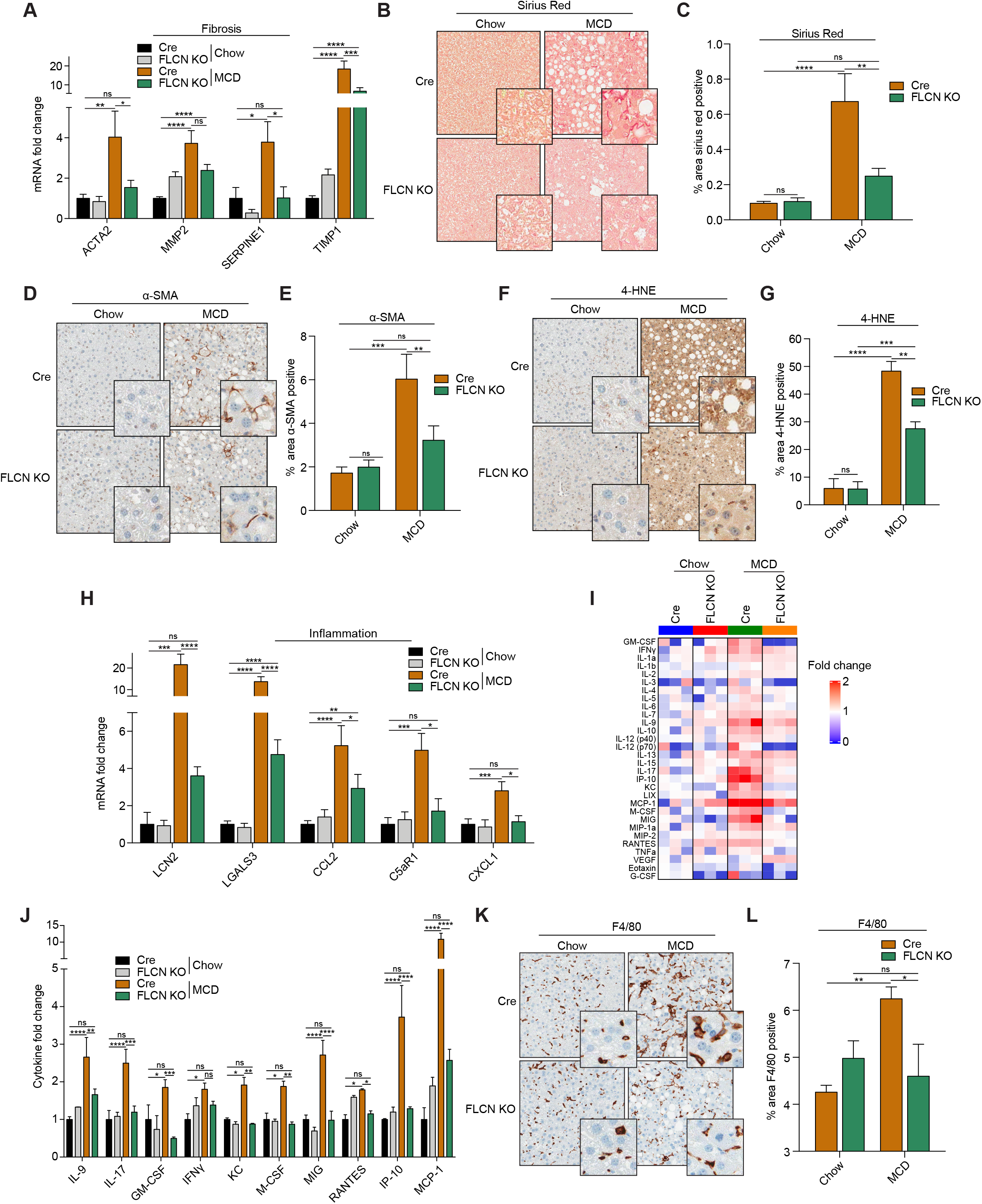
FLCN loss reduces MCD diet induced fibrosis. (**A**) Relative quantitative real-time PCR analysis of fibrosis-related genes mRNA transcript levels in livers of mice fed either on chow or methionine/choline deficient diet (MCD) over a period of 6 weeks (mean ± SEM of the RNA fold change of indicated mRNAs; two-way ANOVA, ns = not significant; *P < 0.05, **P< 0.01, ***P < 0.001, ****P < 0.0001; n = 8 mice per condition). (**B**) Sirius red staining in liver sections of mice fed as described in (A). Images are representative of 6 mice per condition. (**C**) Quantification of the sirius staining in liver sections of mice fed as described in (A) (mean ± SEM, two-way ANOVA, ns = not significant; **P < 0.01; ****P < 0.0001; n = 6 mice per condition) (**D**) α-SMA Immunohistochemistry (IHC) staining in liver sections of mice fed as described in (A). Data are representative of 4 mice per condition. (**E**) Quantification of α-SMA IHC staining in liver sections of mice fed as described in (A) (mean ± SEM, two-way ANOVA, ns = not significant; **P < 0.01; ***P < 0.001; n = 4 mice per condition) (**F**) 4-HNE Immunohistochemistry (IHC) staining in liver sections of mice fed as described in (A). Data are representative of 4 mice per condition. (**G**) Quantification of 4-HNE IHC staining in liver sections of mice fed as described in (A) (mean ± SEM, two-way ANOVA, ns = not significant; **P < 0.01; ***P < 0.001, ****p < 0.0001; n = 4 mice per condition). (**H**) Relative quantitative real-time PCR analysis of inflammation-related genes mRNA transcript levels in livers of mice fed as described in (A) (mean ± SEM of the RNA fold change of indicated mRNAs; two-way ANOVA, ns = not significant; *P < 0.05, **P< 0.01, ****P < 0.0001; n = 8 mice per condition). (**I-J**) Mouse cytokines and chemokines quantification in liver homogenates of mice fed as described in (A) (mean ± SEM, two-way ANOVA, ns = not significant; *P < 0.05, **P< 0.01, ***P < 0.001, ****P < 0.0001; n = 3 mice per condition). (**K**) F4/80 Immunohistochemistry (IHC) staining in liver sections of mice fed as described in (A). Data are representative of 4 mice per condition. (**L**) Quantification of F4/80 IHC staining in liver sections of mice fed as described in (A) (mean ± SEM, two-way ANOVA, ns = not significant; *P < 0.05; **P < 0.01; n = 4 mice per condition)

NASH development is conjointly characterized by increased inflammation that contributes to liver damage and disease progression. In our mouse model, relative mRNA transcript levels of genes involved in inflammation were lower in FLCN KO mice compared to cre mice when fed the MCD diet (Figure 3H). Using a mouse protein cytokine array, we determined the cytokine and chemokine secretion profiles in mouse livers. Cytokine protein levels were higher in cre mice challenged with the MCD diet compared to FLCN KO mice (Figure 3I-J). More specifically, some macrophage chemoattractants (RANTES, MCP-1, IP-10) were induced in cre mice fed the MCD diet but to a lesser extent in FLCN KO mice (Figure 3J). Normal livers are infiltrated by specialized macrophages (Kupffer cells) that contribute to inflammation when activated. Notably, immunohistochemistry staining of F4/80, a marker of murine macrophages, was increased in cre mice fed the MCD diet, but to a lesser extent in FLCN KO mice (Figure 3K, quantified in L). Collectively, our data indicates that liver-specific loss of FLCN reduces inflammation in mice challenged with the MCD diet.

### FLCN loss activates autophagy

FLCN’s best described function to date is involved in autophagy regulation. Loss of FLCN prevents mTORC1 localization on the lysosome surface and subsequently induces autophagy [62–64]. Unsupervised hierarchical clustering of autophagy-related genes from our RNA-seq data revealed an important effect of the diet (cluster 1 and 3, Figure 4A), but also highlighted a subset of genes induced in FLCN KO mice (cluster 2, Figure 4A-B). Relative autophagy-related genes mRNA transcript levels were significantly increased in FLCN KO mice compared to cre mice, while the diet had no effect (Figure 4C). These genes are generally involved in lysosome/autophagosome maturation and activation, which can reduce hepatocellular injury and inflammation by clearing damaged or misfolded proteins and suppressing transcription or maturation of pro-inflammatory cytokines [26–29]. Immunoblot analysis additionally revealed higher protein levels of p62 in cre mice challenged with the MCD diet, indicating impaired autophagy, while FLCN loss reduced p62 protein levels (Figure 4D, quantified in E). Furthermore, the major transcription factors regulating autophagy and lysosome biogenesis, TFE3 and TFEB, were localized in the hepatocyte’s nuclei, depicting again increased autophagy in FLCN KO mice (Figure 4F-I). Taken together, our data suggest that targeting FLCN improves resistance to fibrosis and inflammation while inducing autophagy when challenged with a NASH-inducing diet.

**Figure 4:**
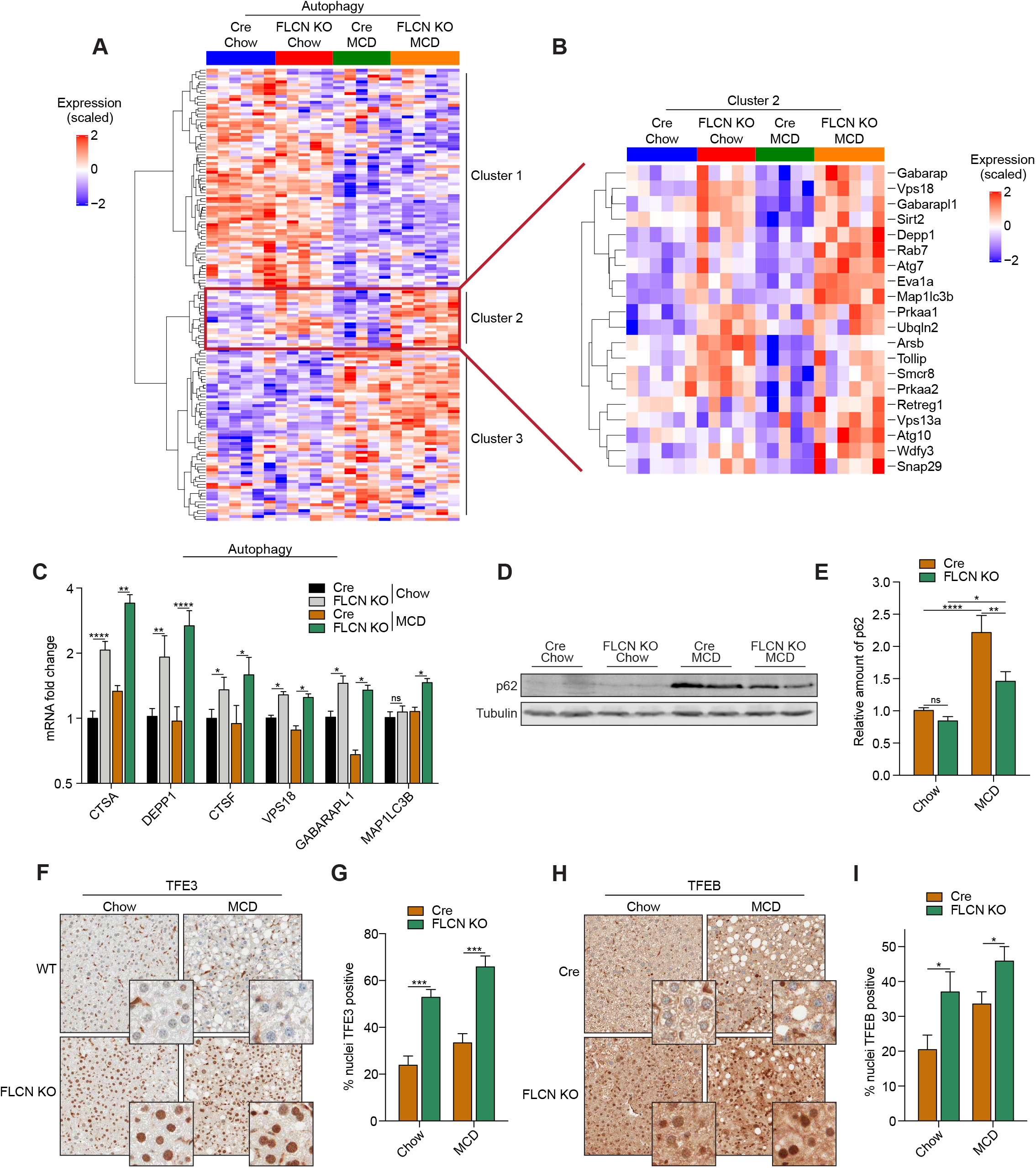
FLCN loss activates autophagy. (**A-B**) Unsupervised hierarchical clustering following RNA-seq analysis in livers of mice fed either on chow or methionine/choline deficient diet (MCD) over a period of 6 weeks. (**C**) Relative quantitative real-time PCR analysis of autophagy-related genes mRNA transcript levels in livers of mice fed as described in (A) (mean ± SEM of the RNA fold change of indicated mRNAs; two-way ANOVA, ns = not significant; *P < 0.05, **P< 0.01, ****P < 0.0001; n = 8 mice per condition). (**D**) Immunoblot of liver protein lysates extracted from mice fed as described in (A). Data are representative of 6 mice per condition. (**E**) Quantification of the relative amount of p62 immunoblot. Data normalized to tubulin levels (mean ± SEM of the relative fold change; two-way ANOVA, ns = not significant; *P < 0.05, **P< 0.01, ****P < 0.0001; n = 6 mice per condition). (**F**) TFE3 Immunohistochemistry (IHC) staining in liver sections of mice fed as described in (A) Data are representative of 6 mice per condition. (**G**) Quantification of TFE3 IHC staining in liver sections of mice fed as described in (A) (mean ± SEM, two-way ANOVA, ***P < 0.001; n = 4 mice per condition). (**H**) TFEB Immunohistochemistry (IHC) staining in liver sections of mice fed as described in (A) Data are representative of 4 mice per condition. (**I**) Quantification of TFEB IHC staining in liver sections of mice fed as described in (A) (mean ± SEM, two-way ANOVA, ***P < 0.001; n = 4 mice per condition).

## Discussion

In the present study, we showed that deletion of *Flcn* in mouse liver tissue reduces fibrosis and inflammation when fed the MCD diet, while activating autophagy. Overall, our results establish a critical role of FLCN in NASH progression, functionally linking autophagy to these diseases. Interestingly, previous groups demonstrated a role for both TFEB and TFE3 in the regulation of lipid metabolism and energy homeostasis in the liver, where TFEB and TFE3 overexpression was sufficient to reverse weight gain and metabolic syndrome in diet-induced and genetic models of obesity, while TFEB/TFE3 deficiency aggravated the phenotype [39,65]. TFEB overexpression also protected from liver damage in alcohol-induced pancreatitis and alcoholic steatohepatitis mouse models [66,67].

Our RNA-seq data revealed major pathways affected by both the diet and FLCN loss. Loss of FLCN specifically in the liver reduced inflammation, fibrosis, and mitotic cell cycle gene expression. The reduction in the cell cycle gene abundance is possibly the consequence of reduced liver damage and therefore of the reduced necessity to regenerate liver cells. Interestingly, others have found that in a model of lung development, FLCN loss impaired WNT signaling in a process completely dependent on TFE3 [68]. This role of FLCN in cell development could explain the lung cyst pathogenesis observed in BHD patients. However, in a different context such as NASH, it is possible that loss of FLCN instead has beneficial effect through similar mechanisms.

We have previously shown that loss of FLCN in *C. elegans* and mammalian cells increased innate immune response in a process dependent on TFEB and TFE3 [43]. Although we observed a decrease in inflammation gene expression in FLCN KO mice fed the MCD diet, we also noted a small increase in gene expression in FLCN KO mice fed on chow compared to Cre (Figure 2E). It is possible that FLCN modulates the immune microenvironment and polarizes the macrophages toward an anti-inflammatory state, which would contribute to the reduced inflammation observed in FLCN KO mice fed on MCD diet. Indeed, it was reported that FLCN loss in hematopoietic stem cells induced CD11c+ CD206+ activated phagocytic macrophages, corresponding to the anti-inflammatory M2 macrophage subtype [69]. However, another study found that bone marrow-derived macrophages from FLCN myeloid KO mice rather promotes spontaneous M1-type polarization and enhanced baseline activation status [47]. Also, RAW 264.7 macrophages targeted with shRNA for FLCN had increased phagocytic activity toward dying/dead cells [70]. The previous effects of FLCN loss on macrophage activation were all dependent on TFEB/TFE3 activation status, highlighting the important role of autophagy in FLCN’s mechanism of regulation. TFEB and TFE3 were also shown to promote efficient autophagy induction in activated macrophages; conversely, TFEB and TFE3 deficient cells had impaired inflammatory response, impaired macrophage differentiation and reduced macrophage infiltration to the site of inflammation [42]. Moreover, TFEB knockdown favored tumor growth with increased infiltration of M2-like macrophages, while TFEB overexpression suppressed the M2 polarization [71]. Therefore, the importance and significance of the basal inflammation observed in FLCN KO livers remains to be investigated.

In conclusion, we revealed that loss of FLCN in liver tissue is involved in autophagy activation, and results in reduction of inflammation and fibrosis. These results show an unexpected role for FLCN in NASH progression. In light of the recently published cryo-EM structures of FLCN [72,73], it is envisioned that pharmacological agents targeting the pocket of the FLCN GAP enzyme could be developed to achieve the loss of function phenotype in the liver shown in this study. Such molecules would represent new possibilities for obesity and NASH treatment through the role of FLCN in hepatocyte homeostasis.

## Material and methods

### Antibodies

The FLCN rabbit polyclonal antibody was generated by the McGill animal resource centre using recombinant GST-FLCN. TFEB (Bethyl Laboratories, cat# A303-673A), TFE3 (Cell Signaling Technology, cat# 14779S), α-SMA (Abcam, cat# 5694), F4/80 (Cell Signaling, cat# 70076), 4-HNE (Millipore, cat# 393207), p62 (Abcam, cat# ab56416), and Tubulin (Sigma, Cat# T9026) antibodies are commercially available.

### Animals

All procedures for generating the *Flcn* knockout mouse model were performed at McGill University. Maintenance and experimental manipulation of mice were performed according to the guidelines and regulations of the McGill University Research and Ethic Animal Committee and the Canadian Council on Animal Care. All studies were carried out using C57BL/6 mice housed on a 12-h light:12-h dark cycle at 22°C. *Flcn*^f/f^ (BHD^f/f^) mice were generated as previously described [74]. To generate liver-specific *Flcn* knockout mice, *Flcn*^f/f^ mice were crossed with Albumin-Cre transgenic mice (kind gift from Dr. André Marette, Laval University, Quebec, Canada). Flcn knockout mice were generated by crossing *Flcn*^f/f^ mice homozygous for the floxed allele with *Flcn*^f/f^; Albumin-Cre^+/-^ mice. Mice were fed on a chow diet containing 10 kcal percent fat (Research Diet, cat# D12450B) or a high fat diet containing 60 kcal percent fat (Research Diet, cat# D12492), or a Methionine/Choline Deficient (MCD) diet (Envigo, cat# TD.90262) beginning at 6 weeks of age and fed ad libidum. Controls used were *Flcn*^WT/WT^; Albumin-cre^+/-^ mice raised in identical conditions. Body weights were measured weekly.

### Protein extraction and immunoblotting

Liver tissues were quickly snap frozen using a BioSpec BioSqueezer (Fisher Scientific, cat# NC1033496) cooled in liquid nitrogen following sacrifice. Approximately 100mg of tissue were solubilized in RIPA buffer using a micropestle and sonicated 10 seconds. The mixture was then centrifugated at 16,000 x *g* for 15 minutes at 4°C and the supernatant collected. Proteins were separated on SDS-PAGE gels and revealed by western blot as previously described [43] using the antibodies listed above and IRDye^®^ 800CW Goat anti-Rabbit IgG Secondary Antibody / IRDye^®^ 680RD Goat anti-Mouse IgG Secondary Antibody (LI-COR Biosciences). Membranes were scanned using the Odyssey imaging system (LI-COR Biosciences) and quantification performed using ImageStudio software.

### Histology

Liver tissues were fixed in 4% formaldehyde for 2 days at room temperature immediately after sacrifice, embedded in paraffin, and cut to 4-μm sections on slides using a Leica Microtome (Leica Biosystems). The slides were stained with hematoxylin and eosin (Leica Biosystems, cat# 3802098), or Sirius Red (Abcam, cat# ab150681) using a Leica ST5020 Multistainer (Leica Biosystems) according to the standard protocol.

### GTTs and ITTs

For the GTTs, mice on either chow or a HFD were subjected to a 16-h fast with free access to water and then injected intraperitoneally with 2 g of 20% D-glucose per kilogram of body weight. The ITTs were performed similarly, with an initial 16 h of fasting and subsequent intraperitoneal injection of 0.75 U of insulin per kilogram of body weight. Blood glucose levels were measured at 0, 15, 30, 60, and 120 min with a OneTouch ultramini glucose meter (OneTouch).

### Serum content quantification

Blood was extracted from cardiac puncture just before sacrifice and incubated with Aprotinin from bovine lung (0.03 TIU, Sigma-Aldrich, cat# A6279) for 2 hours at 4°C. The coagulated blood was spun 10 minutes at 800 x *g* at 4°C and the supernatant (serum) stored at −80°C. Serum total cholesterol, serum free cholesterol, and serum cholesterol esters were quantified using Cholesterol/ Cholesteryl Ester Assay Kit (Abcam, cat# ab65359) according to manufacturer’s instructions. Serum Triglyceride and serum free glycerol were quantified using Serum Triglyceride Determination Kit (Sigma-Aldrich, cat# TR0100) according to manufacturer’s instructions. Serum alanine aminotransferase was quantified using ALT determination kit (Pointe Scientific, cat# A7526) according to manufacturer’s instructions.

### Liver triglyceride quantification

Approximately 10mg of frozen liver tissues were washed with PBS, resuspended in a 2:1 chloroform:methanol solution, and solubilized with a micropestle. The mixture was spun for 10 minutes at 15,000 x *g* at 4°C and the supernatant incubated at 50°C until the chloroform was completely evaporated. The dried pellet was dissolved in a solution containing 60% Butanol, 25% Triton X-100, and 15% Methanol and quantified using Serum Triglyceride Determination Kit (Sigma-Aldrich, cat# TR0100) according to manufacturer’s instructions.

### Quantitative real-time PCR

Total RNA was isolated and purified from approximately 25mg of frozen liver tissue using Total RNA Mini Kit (Geneaid, cat# RT100) according to the manufacturer’s instructions. For quantitative real-time PCR analysis, 0.5 μg of total RNA was reverse-transcribed using the iScript Reverse Transcription Supermix for RT-qPCR (BioRad, cat# 1708841). SYBR Green reactions using the SYBR Green qPCR supermix (BioRad, cat# 1725125) and specific primers (Table 1) were performed using a CFX Connect Real-Time PCR Detection System (BioRad). Relative expression of mRNAs was determined using the software CFX Maestro (BioRad) after normalization against housekeeping genes B2M, TBP, RPLP0, and PTP4a1.

**Table 1:**
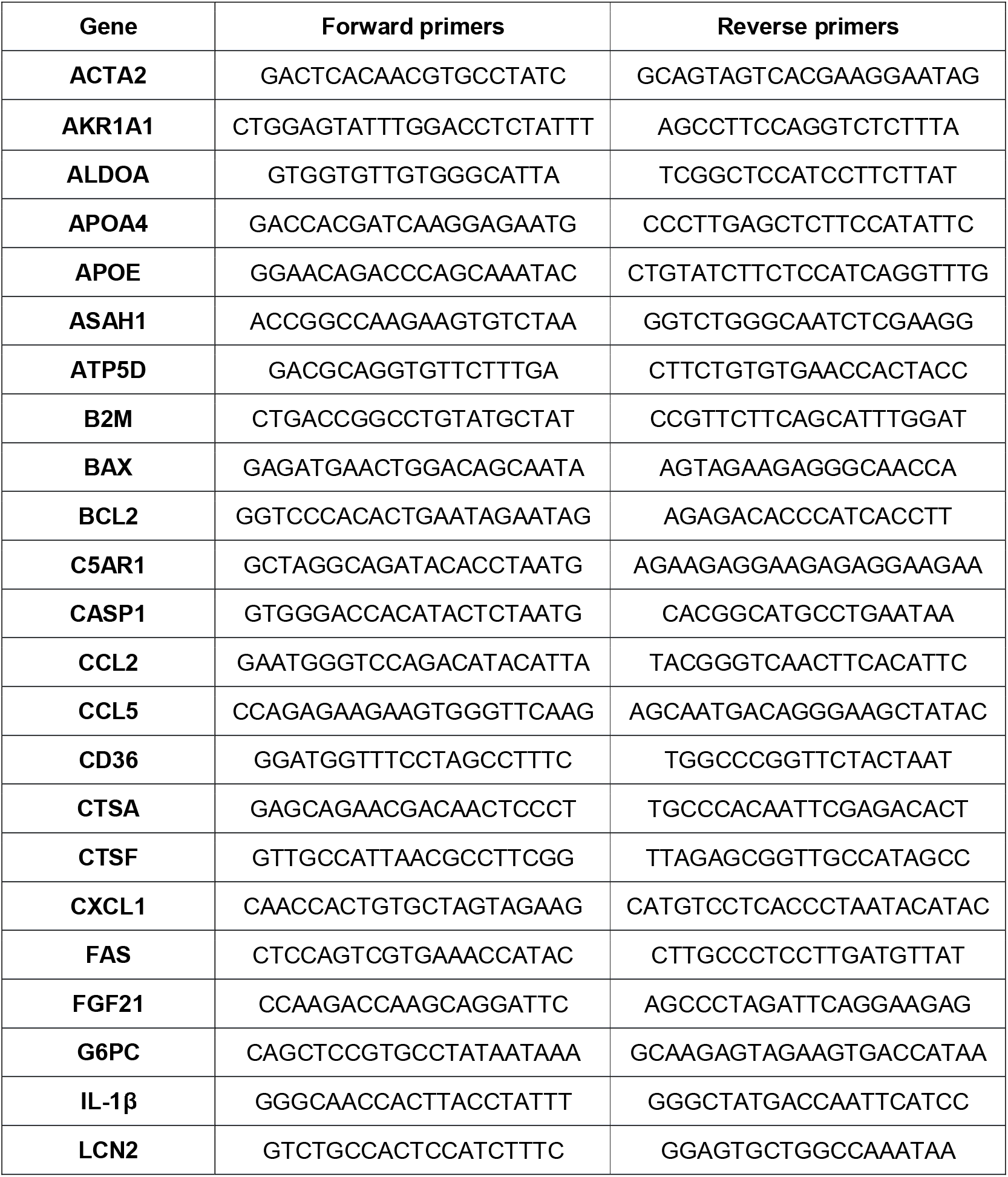
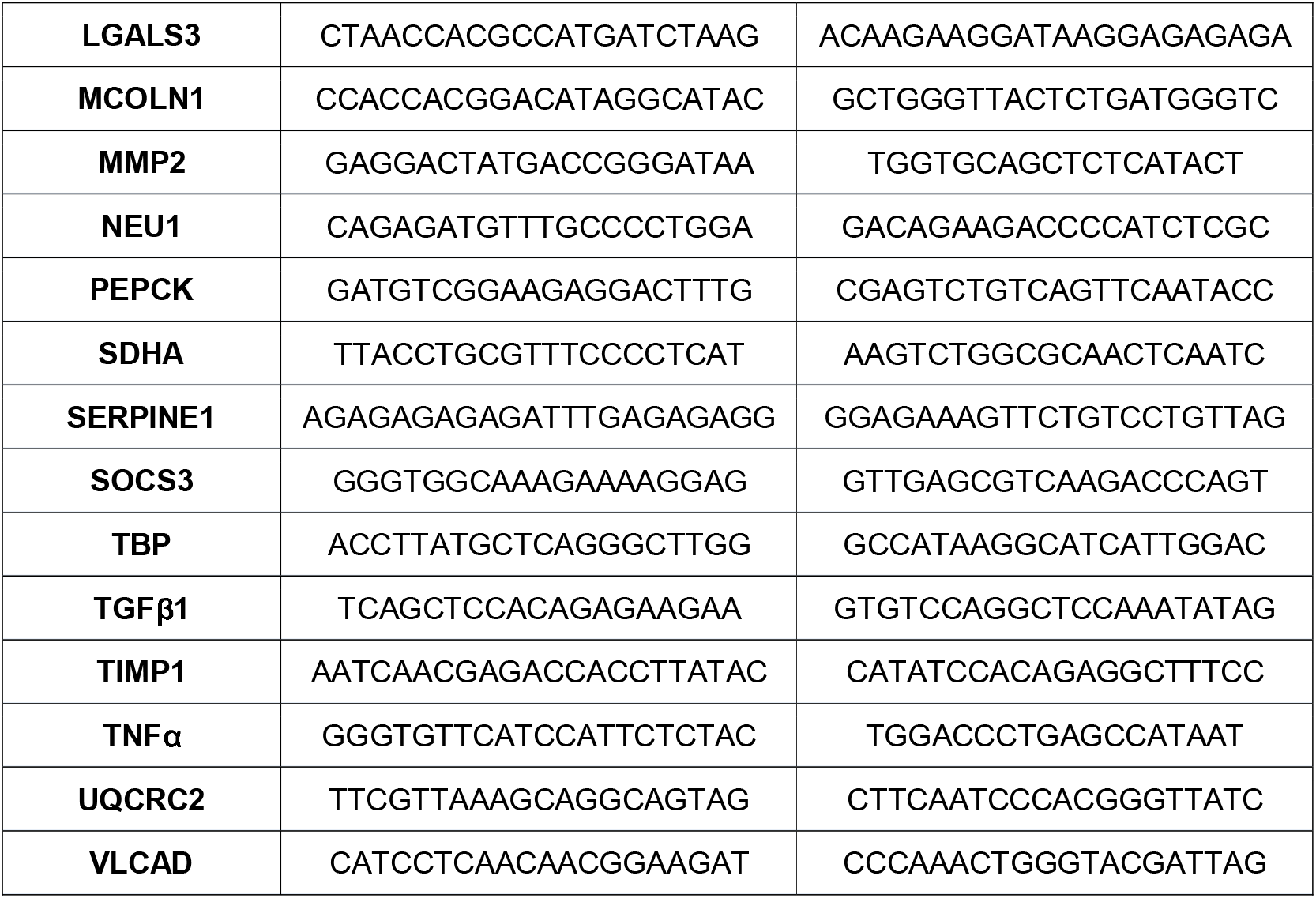
List of mouse primers used in RT-qPCR experiments.

### Immunohistochemistry

Liver tissues embedded in paraffin were cut to 4-μm sections on slides and stained using routine immunohistochemical protocols provided by the GCRC Histology Core using Ventana BenchMark ULTRA system (Roche). Images were acquired using Aperio Scanscope XT (Leica Biosystems) staining quantification was performed using image analysis algorithms in Aperio ImageScope software.

### Mouse Protein Cytokine Array

Approximately 100mg of liver tissue were solubilized in RIPA buffer using a micropestle and sonicated 10 seconds. The mixture was then centrifugated at 16,000 x g for 15 minutes at 4°C and the supernatant collected. 31 mouse cytokine/chemokine biomarkers were simultaneously quantified by using a Discovery Assay^®^ called the Mouse Cytokine Array / Chemokine Array 31-Plex (Eve Technologies Corp, cat# MD31)

### RNA-seq analysis

Total RNA was isolated and purified from approximately 25mg of frozen liver tissue using Total RNA Mini Kit (Geneaid, cat# RT100) according to the manufacturer’s instructions. Library preparation and sequencing was made at the Institute for Research in Immunology and Cancer’s Genomics Platform (IRIC). 500 ng of total RNA was used for library preparation. RNA quality control was assessed with the Bioanalyzer RNA 6000 Nano assay on the 2100 Bioanalyzer system (Agilent technologies). Library preparation was done with the KAPA mRNAseq Hyperprep kit (KAPA, cat# KK8581). Ligation was made with Illumina dual-index UMI (IDT) and 10 PCR cycles was required to amplify cDNA libraries. Libraries were quantified by QuBit and BioAnalyzer DNA1000. All libraries were diluted to 10 nM and normalized by qPCR using the KAPA library quantification kit (KAPA, cat# KK4973). Libraries were pooled to equimolar concentration. Sequencing was performed with the Illumina Nextseq500 using the Nextseq High Output 75 (1×75bp) cycles kit using 2.6 pM of the pooled libraries. Around 25 M single-end PF reads were generated per sample.

Following data acquisition, adaptor sequences and low quality score bases (Phred score < 30) were first trimmed using Trimmomatic [75]. The resulting reads were aligned to the GRCm38 mouse reference genome assembly, using STAR [76]. Read counts are obtained using HTSeq [77] and are represented as a table, which reports for each sample (columns), the number of reads mapped to a given gene (rows). For all downstream analyses, we excluded lowly-expressed genes with an average read count lower than 10 across all samples, resulting in 14,057 genes in total. The R package limma [78] was used to identify differences in gene expression levels between the different conditions. Nominal p-values were corrected for multiple testing using the Benjamini-Hochberg method. To assess the effect FLCN-KO in the response to MCD-diet, we first obtained DE genes (FDR < 0.05 and |log2FC| > 1) in Cre.MCD_diet vs Cre.Chow and Flcn-KO.MCD_diet vs Flcn-KO.Chow, and then filtered for those that show |difference in log2FC| > 1 (differentially responsive (DR) genes). Unsupervised hierarchical clustering of the 1,110 DR genes shows three distinct patterns of changes in expression. Pathway enrichment analyses were performed using Enrichr [79].

### Statistical Analyses

Data are expressed as mean ±SEM. All experiments and measurements were performed on at least 3 mice as indicated. Statistical analyses for all data were performed using student’s t-test, one-way ANOVA or two-way ANOVA as indicated using GraphPad Prism 7 software. Statistical significance is indicated in figures (*P<0.05, **P<0.01, ***P<0.001, ****P<0.0001).

### Data availability

RNA-sequencing data has been deposited in the Gene Expression Omnibus under the accession GSE156918.

## Acknowledgements

M.P., L.E-H., J.M.J.R.R. were supported by CIHR, FRQS, and CONACYT, respectively. This work was supported by grants to A.P. from the Kidney Foundation of Canada, Terry Fox foundation (TFF-166128), CIHR (PJT-165829) and the Cancer Research Society. We would like to thank Hannah Zhang, the Canadian Centre for Computational Genomics, a Genome Canada funded bioinformatics platform, and the McGill University Goodman Cancer Research Centre Histology Core Facility for their support and advices.

## Author contributions

Conception and design of the experiments: M.P., L.EH., M.Y., J.L.E., P.M.S., and A.P.; collection, assembly, analysis and interpretation of data: M.P., L.EH., M.B., M.Y., J.M.J.R.R., and H.J.; drafting the article or revising it critically for important intellectual content: M.P., L.EH., M.B., M.Y., J.M.J.R.R. H.J., J.L.E., P.M.S., and A.P.

---

**Supplemental figure 1: FLCN loss reduces high fat diet induced NAFLD**

(**A**) Immunoblot of protein lysates extracted from livers of 5-months old Alb-cre WT (Cre) and Alb-cre FLCN KO (FLCN KO) mice. Results are representative of 6 mice per condition. (**B**) Schematic showing the generation of liver-specific KO mice. Mice carrying Flcn alleles flanked by loxP sites were crossed with Albumin-Cre transgenic mice to generate liver-specific Flcn knockout mice. (**C**) Schematic showing FLCN domains aligned on the corresponding exons on the mRNA sequence. Exon 7 is removed following expression of Albumin-Cre. (**D**) Mouse body weight when fed either on chow or high fat diet (HFD) over a period of 6 weeks (mean ± SEM, two-way ANOVA, *P < 0.05; **P <0.01; ****P < 0.0001 compared to Cre-chow, ^#^P < 0.05; ^##^P < 0.01; ^####^P < 0.0001 compared to Cre-HFD; n = 6 mice per condition). (**E**) Representative images of H&E staining in liver sections of mice fed either chow or HFD over a period of 6 weeks. Images are representative of 6 mice per condition. (**F**) Blood glucose during a GTT in chow or HFD-fed mice following a 16-h fast and intraperitoneal glucose administration of 2 g per kg of body weight (mean ± SEM, two-way ANOVA, *P < 0.05; ***P <0.001 compared to Cre-chow, #P < 0.05 compared to Cre-HFD; n = 6 mice per condition). (**G**) Blood glucose during an ITT in chow or HFD-fed mice following a 16-h fast and intraperitoneal insulin administration of 0.75 U per kg of body weight. (mean ± SEM, two-way ANOVA, **P < 0.01; ***P <0.001 compared to Cre-chow, #P < 0.05 compared to Cre-HFD; n = 6 mice per condition).

**Supplemental figure 2: Weight and blood parameters are unaffected by FLCN loss**

(**A**) Mouse body weight when fed either on chow or methionine/choline deficient diet (MCD) over a 6-week period in males (left) and females (right) (mean ± SEM, two-way ANOVA, ns = not significant; ****P < 0.0001; n = 8 mice per condition). (**B**) Serum total cholesterol quantification in male (left) and female (right) mice fed as described in (A) (mean ± SEM, two-way ANOVA, ns = not significant; ****P < 0.0001; n = 8 mice per condition). (**C**) Serum free cholesterol quantification in male (left) and female (right) mice fed as described in (A) (mean ± SEM, two-way ANOVA, ns = not significant; ****P < 0.0001; n = 8 mice per condition). (**D**) Serum cholesterol esters quantification in male (left) and female (right) mice fed as described in (A) (mean ± SEM, two-way ANOVA, ns = not significant; ****P < 0.0001; n = 8 mice per condition).

**Supplemental figure 3: Gene expression analysis in liver tissues of Cre and FLCN KO mice** (**A-E**) Unsupervised hierarchical clustering following RNA-seq analysis of genes related to apoptosis (A, GO:0006915), lipolysis (B, GO:0016042), lipogenesis (C, GO:0008610), glycolysis (D, GO:0006096), and gluconeogenesis (E, GO:0006094) in livers of mice fed either chow or methionine/choline deficient diet (MCD) over a period of 6 weeks. (**B**) Relative quantitative real-time PCR analysis in livers of mice fed as described in (A) (mean ± SEM of the RNA fold change of indicated mRNAs; two-way ANOVA, ns = not significant; *P< 0.05; **P< 0.01; n = 8 mice per condition).

## Abbreviations

FLCN: Folliculin
HFD: High fat diet
H&E: Hematoxylin and eosin
MCD: Methionine/Choline Deficient diet
mTORC1: mammalian Target of Rapamycin Complex 1
NAFLD: Non-alcoholic fatty liver disease
NASH: Non-alcoholic steatohepatitis
TFEB: Transcription Factor EB
TFE3: Transcription Factor E3

## References

1. Singh GM, Danaei G, Farzadfar F, et al. The age-specific quantitative effects of metabolic risk factors on cardiovascular diseases and diabetes: a pooled analysis. PLoS One. 2013;8:e65174.

2. Czernichow S, Kengne AP, Stamatakis E, et al. Body mass index, waist circumference and waist-hip ratio: which is the better discriminator of cardiovascular disease mortality risk?: evidence from an individual-participant meta-analysis of 82 864 participants from nine cohort studies. Obes Rev. 2011;12:680–687.

3. Anstey KJ, Cherbuin N, Budge M, et al. Body mass index in midlife and late-life as a risk factor for dementia: a meta-analysis of prospective studies. Obes Rev. 2011;12:e426–37.

4. Lauby-Secretan B, Scoccianti C, Loomis D, et al. Body Fatness and Cancer--Viewpoint of the IARC Working Group. N Engl J Med. 2016;375:794–798.

5. Chooi YC, Ding C, Magkos F. The epidemiology of obesity. Metab Clin Exp. 2019;92:6–10.

6. European Association for the Study of the Liver (EASL), European Association for the Study of Diabetes (EASD), European Association for the Study of Obesity (EASO). EASL-EASD-EASO Clinical Practice Guidelines for the Management of Non-Alcoholic Fatty Liver Disease. Obes Facts. 2016;9:65–90.

7. Arab JP, Arrese M, Trauner M. Recent Insights into the Pathogenesis of Nonalcoholic Fatty Liver Disease. Annu Rev Pathol. 2018;13:321–350.

8. Satapathy SK, Sanyal AJ. Epidemiology and natural history of nonalcoholic fatty liver disease. Semin Liver Dis. 2015;35:221–235.

9. Angulo P, Kleiner DE, Dam-Larsen S, et al. Liver Fibrosis, but No Other Histologic Features, Is Associated With Long-term Outcomes of Patients With Nonalcoholic Fatty Liver Disease. Gastroenterology. 2015;149:389–97.e10.

10. Roeb E, Geier A. Nonalcoholic steatohepatitis (NASH) - current treatment recommendations and future developments. Z Gastroenterol. 2019;57:508–517.

11. Younossi ZM. Non-alcoholic fatty liver disease - A global public health perspective. J Hepatol. 2019;70:531–544.

12. Ibrahim SH, Hirsova P, Gores GJ. Non-alcoholic steatohepatitis pathogenesis: sublethal hepatocyte injury as a driver of liver inflammation. Gut. 2018;67:963–972.

13. Serviddio G, Bellanti F, Tamborra R, et al. Alterations of hepatic ATP homeostasis and respiratory chain during development of non-alcoholic steatohepatitis in a rodent model. Eur J Clin Invest. 2008;38:245–252.

14. Begriche K, Massart J, Robin M-A, et al. Mitochondrial adaptations and dysfunctions in nonalcoholic fatty liver disease. Hepatology. 2013;58:1497–1507.

15. Machado MV, Diehl AM. Pathogenesis of nonalcoholic steatohepatitis. Gastroenterology. 2016;150:1769–1777.

16. Ashraf NU, Sheikh TA. Endoplasmic reticulum stress and Oxidative stress in the pathogenesis of Non-alcoholic fatty liver disease. Free Radic Res. 2015;49:1405–1418.

17. Agosti P, Sabbà C, Mazzocca A. Emerging metabolic risk factors in hepatocellular carcinoma and their influence on the liver microenvironment. Biochim Biophys Acta Mol Basis Dis. 2018;1864:607–617.

18. Koo S-Y, Park E-J, Lee C-W. Immunological distinctions between nonalcoholic steatohepatitis and hepatocellular carcinoma. Exp Mol Med. 2020;

19. Angulo P, Machado MV, Diehl AM. Fibrosis in nonalcoholic Fatty liver disease: mechanisms and clinical implications. Semin Liver Dis. 2015;35:132–145.

20. Machado MV, Diehl AM. Liver renewal: detecting misrepair and optimizing regeneration. Mayo Clin Proc. 2014;89:120–130.

21. González-Rodríguez A, Mayoral R, Agra N, et al. Impaired autophagic flux is associated with increased endoplasmic reticulum stress during the development of NAFLD. Cell Death Dis. 2014;5:e1179.

22. Inami Y, Yamashina S, Izumi K, et al. Hepatic steatosis inhibits autophagic proteolysis via impairment of autophagosomal acidification and cathepsin expression. Biochem Biophys Res Commun. 2011;412:618–625.

23. Koga H, Kaushik S, Cuervo AM. Altered lipid content inhibits autophagic vesicular fusion. FASEB J. 2010;24:3052–3065.

24. Yang L, Li P, Fu S, et al. Defective hepatic autophagy in obesity promotes ER stress and causes insulin resistance. Cell Metab. 2010;11:467–478.

25. Lavallard VJ, Gual P. Autophagy and non-alcoholic fatty liver disease. Biomed Res Int. 2014;2014:120179.

26. Wang Y, Singh R, Xiang Y, et al. Macroautophagy and chaperone-mediated autophagy are required for hepatocyte resistance to oxidant stress. Hepatology. 2010;52:266–277.

27. Lemasters JJ. Selective mitochondrial autophagy, or mitophagy, as a targeted defense against oxidative stress, mitochondrial dysfunction, and aging. Rejuvenation Res. 2005;8:3–5.

28. Lee H-M, Shin D-M, Yuk J-M, et al. Autophagy negatively regulates keratinocyte inflammatory responses via scaffolding protein p62/SQSTM1. J Immunol. 2011;186:1248–1258.

29. Saitoh T, Fujita N, Jang MH, et al. Loss of the autophagy protein Atg16L1 enhances endotoxin-induced IL-1beta production. Nature. 2008;456:264–268.

30. Sardiello M, Palmieri M, di Ronza A, et al. A gene network regulating lysosomal biogenesis and function. Science. 2009;325:473–477.

31. Paquette M, El-Houjeiri L, Pause A. mTOR Pathways in Cancer and Autophagy. Cancers (Basel). 2018;10.

32. Shimobayashi M, Hall MN. Making new contacts: the mTOR network in metabolism and signalling crosstalk. Nat Rev Mol Cell Biol. 2014;15:155–162.

33. Martina JA, Chen Y, Gucek M, et al. MTORC1 functions as a transcriptional regulator of autophagy by preventing nuclear transport of TFEB. Autophagy. 2012;8:903–914.

34. Roczniak-Ferguson A, Petit CS, Froehlich F, et al. The transcription factor TFEB links mTORC1 signaling to transcriptional control of lysosome homeostasis. Sci Signal. 2012;5:ra42.

35. Settembre C, Zoncu R, Medina DL, et al. A lysosome-to-nucleus signalling mechanism senses and regulates the lysosome via mTOR and TFEB. EMBO J. 2012;31:1095–1108.

36. Vega-Rubin-de-Celis S, Peña-Llopis S, Konda M, et al. Multistep regulation of TFEB by MTORC1. Autophagy. 2017;13:464–472.

37. Martina JA, Diab HI, Lishu L, et al. The nutrient-responsive transcription factor TFE3 promotes autophagy, lysosomal biogenesis, and clearance of cellular debris. Sci Signal. 2014;7:ra9.

38. Palmieri M, Impey S, Kang H, et al. Characterization of the CLEAR network reveals an integrated control of cellular clearance pathways. Hum Mol Genet. 2011;20:3852–3866.

39. Settembre C, De Cegli R, Mansueto G, et al. TFEB controls cellular lipid metabolism through a starvation-induced autoregulatory loop. Nat Cell Biol. 2013;15:647–658.

40. Settembre C, Di Malta C, Polito VA, et al. TFEB links autophagy to lysosomal biogenesis. Science. 2011;332:1429–1433.

41. Najibi M, Labed SA, Visvikis O, et al. An Evolutionarily Conserved PLC-PKD-TFEB Pathway for Host Defense. Cell Rep. 2016;15:1728–1742.

42. Pastore N, Brady OA, Diab HI, et al. TFEB and TFE3 cooperate in the regulation of the innate immune response in activated macrophages. Autophagy. 2016;12:1240–1258.

43. El-Houjeiri L, Possik E, Vijayaraghavan T, et al. The Transcription Factors TFEB and TFE3 Link the FLCN-AMPK Signaling Axis to Innate Immune Response and Pathogen Resistance. Cell Rep. 2019;26:3613–3628.e6.

44. Martina JA, Puertollano R. Rag GTPases mediate amino acid-dependent recruitment of TFEB and MITF to lysosomes. J Cell Biol. 2013;200:475–491.

45. Hong S-B, Oh H, Valera VA, et al. Inactivation of the FLCN tumor suppressor gene induces TFE3 transcriptional activity by increasing its nuclear localization. PLoS One. 2010;5:e15793.

46. Wada S, Neinast M, Jang C, et al. The tumor suppressor FLCN mediates an alternate mTOR pathway to regulate browning of adipose tissue. Genes Dev. 2016;30:2551–2564.

47. Li J, Wada S, Weaver LK, et al. Myeloid Folliculin balances mTOR activation to maintain innate immunity homeostasis. JCI Insight. 2019;5.

48. Schmidt LS, Linehan WM. FLCN: The causative gene for Birt-Hogg-Dubé syndrome. Gene. 2018;640:28–42.

49. Sancak Y, Bar-Peled L, Zoncu R, et al. Ragulator-Rag complex targets mTORC1 to the lysosomal surface and is necessary for its activation by amino acids. Cell. 2010;141:290–303.

50. Tsun Z-Y, Bar-Peled L, Chantranupong L, et al. The folliculin tumor suppressor is a GAP for the RagC/D GTPases that signal amino acid levels to mTORC1. Mol Cell. 2013;52:495–505.

51. Péli-Gulli M-P, Sardu A, Panchaud N, et al. Amino Acids Stimulate TORC1 through Lst4-Lst7, a GTPase-Activating Protein Complex for the Rag Family GTPase Gtr2. Cell Rep. 2015;13:1–7.

52. Petit CS, Roczniak-Ferguson A, Ferguson SM. Recruitment of folliculin to lysosomes supports the amino acid-dependent activation of Rag GTPases. J Cell Biol. 2013;202:1107–1122.

53. Possik E, Jalali Z, Nouët Y, et al. Folliculin regulates ampk-dependent autophagy and metabolic stress survival. PLoS Genet. 2014;10:e1004273.

54. Possik E, Ajisebutu A, Manteghi S, et al. FLCN and AMPK confer resistance to hyperosmotic stress via remodeling of glycogen stores. PLoS Genet. 2015;11:e1005520.

55. Hasumi H, Baba M, Hasumi Y, et al. Regulation of mitochondrial oxidative metabolism by tumor suppressor FLCN. J Natl Cancer Inst. 2012;104:1750–1764.

56. Yan M, Audet-Walsh É, Manteghi S, et al. Chronic AMPK activation via loss of FLCN induces functional beige adipose tissue through PGC-1α/ERRα. Genes Dev. 2016;30:1034–1046.

57. Postic C, Shiota M, Niswender KD, et al. Dual roles for glucokinase in glucose homeostasis as determined by liver and pancreatic beta cell-specific gene knock-outs using Cre recombinase. J Biol Chem. 1999;274:305–315.

58. Van Herck MA, Vonghia L, Francque SM. Animal Models of Nonalcoholic Fatty Liver Disease-A Starter’s Guide. Nutrients. 2017;9.

59. Itagaki H, Shimizu K, Morikawa S, et al. Morphological and functional characterization of non-alcoholic fatty liver disease induced by a methionine-choline-deficient diet in C57BL/6 mice. Int J Clin Exp Pathol. 2013;6:2683–2696.

60. Caldez MJ, Bjorklund M, Kaldis P. Cell cycle regulation in NAFLD: when imbalanced metabolism limits cell division. Hepatol Int. 2020;14:463–474.

61. Burt AD, Lackner C, Tiniakos DG. Diagnosis and assessment of NAFLD: definitions and histopathological classification. Semin Liver Dis. 2015;35:207–220.

62. Meng J, Ferguson SM. GATOR1-dependent recruitment of FLCN-FNIP to lysosomes coordinates Rag GTPase heterodimer nucleotide status in response to amino acids. J Cell Biol. 2018;217:2765–2776.

63. Hosios AM, Manning BD. Lysosomal catch-and-release controls mTORC1. Nat Cell Biol. 2018;20:996–997.

64. Shen K, Huang RK, Brignole EJ, et al. Architecture of the human GATOR1 and GATOR1-Rag GTPases complexes. Nature. 2018;556:64–69.

65. Pastore N, Vainshtein A, Klisch TJ, et al. TFE3 regulates whole-body energy metabolism in cooperation with TFEB. EMBO Mol Med. 2017;9:605–621.

66. Chao X, Wang S, Zhao K, et al. Impaired TFEB-Mediated Lysosome Biogenesis and Autophagy Promote Chronic Ethanol-Induced Liver Injury and Steatosis in Mice. Gastroenterology. 2018;155:865–879.e12.

67. Wang S, Ni H-M, Chao X, et al. Critical Role of TFEB-Mediated Lysosomal Biogenesis in Alcohol-Induced Pancreatitis in Mice and Humans. Cell Mol Gastroenterol Hepatol. 2020;10:59–81.

68. Kennedy JC, Khabibullin D, Hougard T, et al. Loss of FLCN inhibits canonical WNT signaling via TFE3. Hum Mol Genet. 2019;28:3270–3281.

69. Endo M, Baba M, Endoh T, et al. The FLCN-TFE3 axis regulates macrophage activation through cellular metabolism. Exp Hematol. 2017;53:S108–S109.

70. Endoh M, Baba M, Endoh T, et al. A FLCN-TFE3 Feedback Loop Prevents Excessive Glycogenesis and Phagocyte Activation by Regulating Lysosome Activity. Cell Rep. 2020;30:1823–1834.e5.

71. Fang L, Hodge J, Saaoud F, et al. Transcriptional factor EB regulates macrophage polarization in the tumor microenvironment. Oncoimmunology. 2017;6:e1312042.

72. Shen K, Rogala KB, Chou H-T, et al. Cryo-EM Structure of the Human FLCN-FNIP2-Rag-Ragulator Complex. Cell. 2019;179:1319–1329.e8.

73. Lawrence RE, Fromm SA, Fu Y, et al. Structural mechanism of a Rag GTPase activation checkpoint by the lysosomal folliculin complex. Science. 2019;366:971–977.

74. Baba M, Furihata M, Hong S-B, et al. Kidney-targeted Birt-Hogg-Dube gene inactivation in a mouse model: Erk1/2 and Akt-mTOR activation, cell hyperproliferation, and polycystic kidneys. J Natl Cancer Inst. 2008;100:140–154.

75. Bolger AM, Lohse M, Usadel B. Trimmomatic: a flexible trimmer for Illumina sequence data. Bioinformatics. 2014;30:2114–2120.

76. Dobin A, Davis CA, Schlesinger F, et al. STAR: ultrafast universal RNA-seq aligner. Bioinformatics. 2013;29:15–21.

77. Anders S, Pyl PT, Huber W. HTSeq — a Python framework to work with high-throughput sequencing data. Bioinformatics. 2015;31:166–169.

78. Ritchie ME, Phipson B, Wu D, et al. limma powers differential expression analyses for RNA-sequencing and microarray studies. Nucleic Acids Res. 2015;43:e47.

79. Kuleshov MV, Jones MR, Rouillard AD, et al. Enrichr: a comprehensive gene set enrichment analysis web server 2016 update. Nucleic Acids Res. 2016;44:W90–7.

