## Supplementary figures and images for "*FLCN* Gene Ablation Reduces Fibrosis and Inflammation in a Diet-Induced NASH Model"

### Supplemental figures

#### Supplemental Figure 1

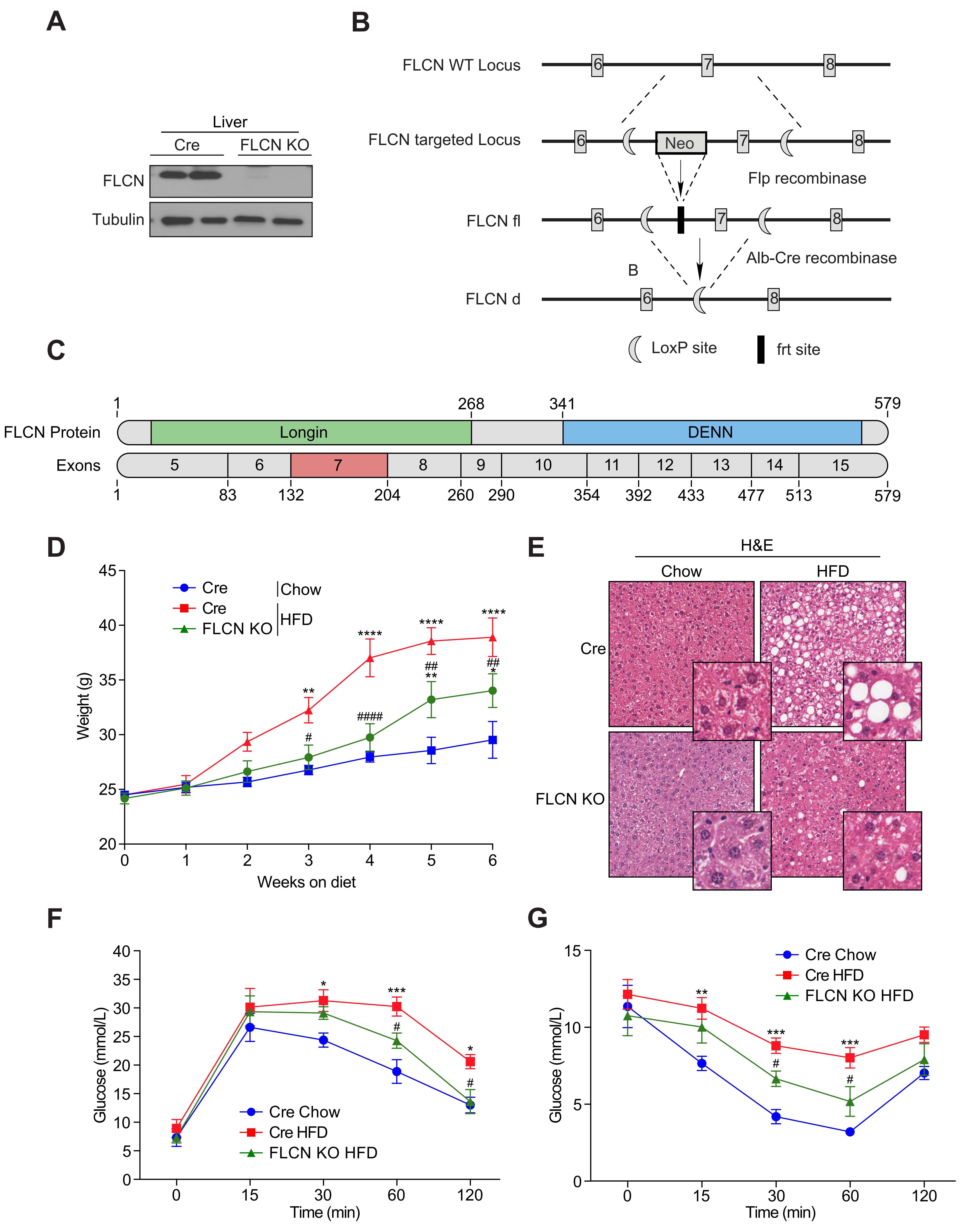

#### Supplemental Figure 2

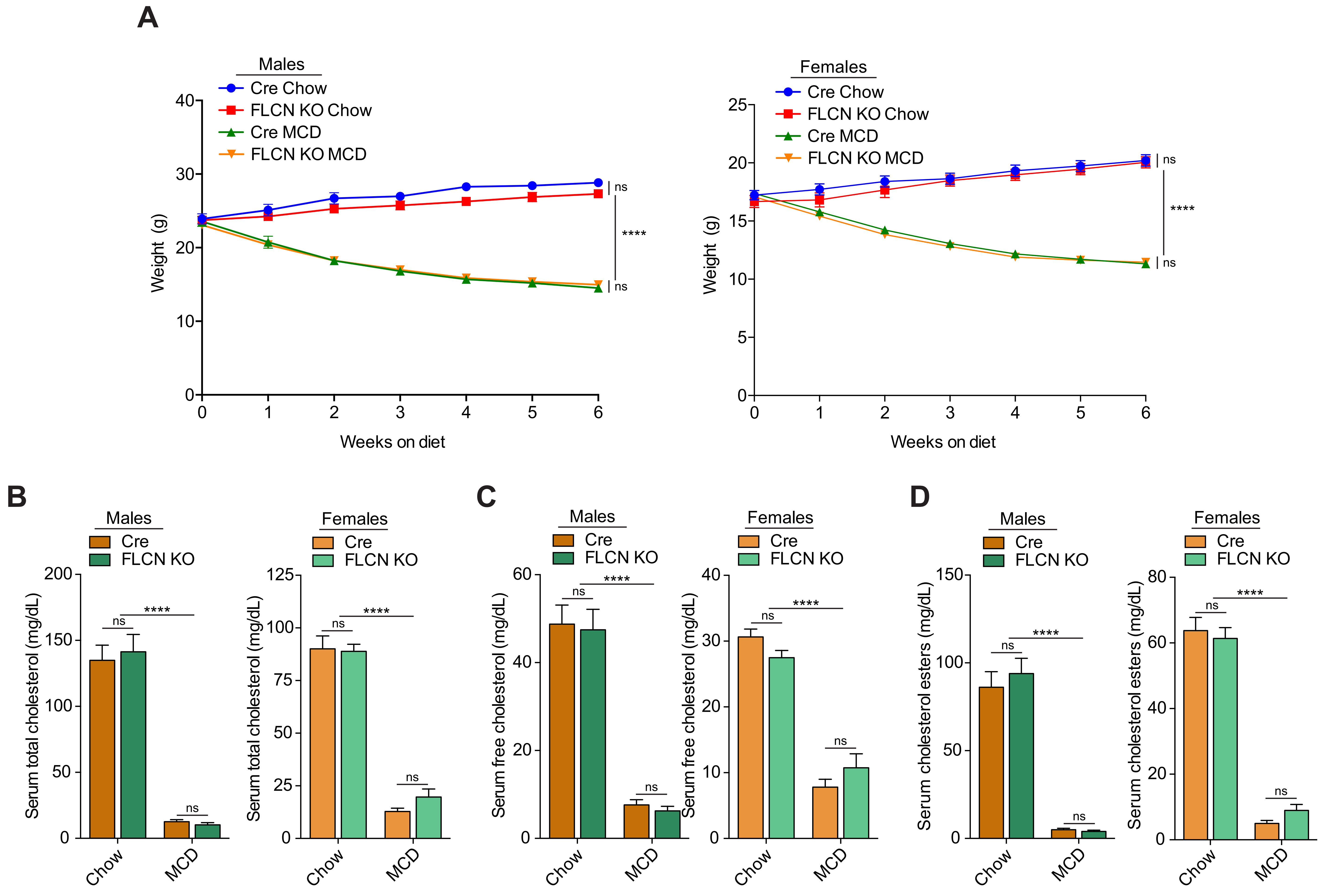

##### Supplemental Figure 3

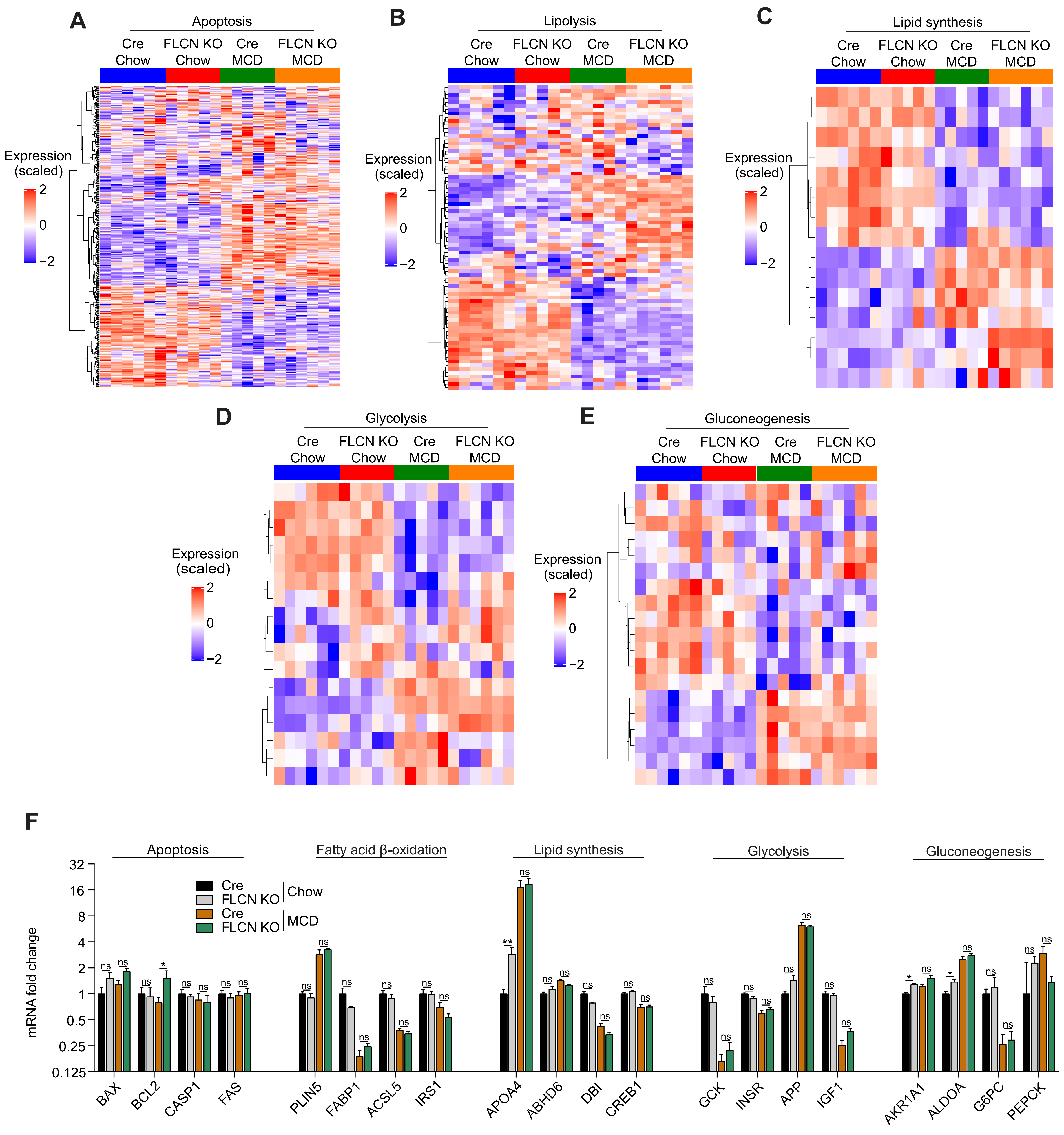
